# Characterization of the novel *SmsHSP24.1* Promoter: Unveiling its abiotic stress-inducible expression in transgenic Eggplant (*Solanum melongena* L.)

**DOI:** 10.1101/2024.04.20.589358

**Authors:** Mst. Muslima Khatun, Imran Khan, Bhabesh Borphukan, Keshob Chandra Das, Haseena Khan, Mohammad Riazul Islam, Malireddy K Reddy, Md Salimullah

## Abstract

Promoters play a pivotal role in regulating gene expression, orchestrating vital processes in plants, including their responses to various environmental stresses. In this study, we focus on the comprehensive characterization of the *SmsHSP24.1* promoter, a novel *cis*-acting element, within the context of transgenic Eggplant (*Solanum melongena* L.). This detailed analysis shed light on the intricate mechanisms governing *SmsHSP24.1* promoter-driven gene regulation, particularly in response to adverse environmental challenges such as heat, salt and drought stressors offering valuable insights into its role in plant stress adaptation. The advances in our understanding of promoter-driven gene regulation also contribute to the broader goal of enhancing crop resilience to abiotic stresses, positioning the *SmsHSP24.1* promoter as a promising tool in agricultural biotechnology applications.

**Highlights:**

1. Demonstrated that the full-length 2.0 kb SmsHSP24.1 promoter significantly enhances gene expression under heat stress, with an observable decline in expression with promoter truncation.
2. Identified specific regulatory elements within the SmsHSP24.1 promoter that are decisive for inducible expression in response to abiotic stresses such as heat, salt, and drought.
3. Highlighted the utility of the SmsHSP24.1 promoter in crop improvement programs, offering a tool for developing transgenic plant tolerance to combined stresses.

## 1. Introduction

Plant growth, development, and adaptation are intricately regulated by gene expression. Central to this regulatory process are promoters, and cis-acting elements that initiate transcription and serve as key determinants of gene expression patterns. While constitutive promoters are commonly employed for gene characterisation and transgenic studies in plants, their inherent lack of specificity and limited control over protein expression pose significant limitations. Studies suggest that ectopic expression of an exogenous gene under the control of a constitutive promoter often results in unexpected phenotypic alterations, such as reduced plant growth or premature or early flowering and yield penalty (Jeong & Jung, 2015; Kummari et al., 2020; Misra & Ganesan, 2021). This lack of precision in gene regulation has prompted the exploration of inducible promoters that offer more targeted control over transgene expression in response to specific plant tissue growth and development cues or environmental factors. Thus, inducible promoters offer the advantage of precise spatiotemporal control over transgene expression, facilitating targeted investigations into gene function under specific conditions or tissue specific study with minimal defect on growth and development (Kummari et al., 2020). With a comprehensive understanding of plant inducible promoters, boasting precise spatiotemporal control, the potential applications extend even to harnessing the precision of the CRISPR/Cas9 gene-editing toolbox. This enables targeted overexpression or knockdown of genes associated with pivotal agroclimatic traits. Recent studies demonstrated a comparative analysis of using CRISPR/Cas9 system under a heat shock inducible promoter produces less off-target mutations compared to constitutive ubiquitin promoter in Arabidopsis, rice, and tobacco (Liang et al., 2023; Nandy et al., 2019; Sheva et al., 2020).

Thus, this emerging approach represents a novel plant breeding toolbox poised to revolutionize crop improvement through precise gene manipulation. Hence, the pursuit of inducible promoters for precise spatiotemporal gene control is paramount. The cloning and characterization of diverse inducible promoters tailored for plant genetic transformation have become an imperative undertaking. Notably, in the realm of both transgenic applications and CRISPR toolbox-based plant gene editing, a spectrum of inducible promoters has been elucidated. These promoters can be activated by specific chemicals (Cao et al., 2001; Darwish et al., 2014; Khan et al., 2011) or stimuli, including but not limited to heat (Freeman et al., 2011; Harrington et al., 2020), cold (Khodakovskaya et al., 2006; Li et al., 2013; Tittarelli et al., 2009), drought (Estrada-Melo et al., 2015; Sun et al., 2024), mechanical injury (Pandey et al., 2019; Yan et al., 2013), light (Gligorovski et al., 2023; Timerbaev & Dolgov, 2019), pathogen (Heise et al., 2002; Kluge et al., 2018) and salt (Peethambaran et al., 2018; Yan et al., 2023). This research focuses on the comprehensive characterization of the SmsHSP24.1 promoter and its role in responding to abiotic stress factors in Eggplant (Solanum melongena). In our prior investigations, we have demonstrated the remarkable sensitivity of the SmsHSP24.1 protein to a spectrum of environmental stressors, including heat, salt, and mannitol (Khatun et al., 2021). This heightened transcript expression in response to stress signifies the pivotal role played by SmsHSP24.1 in the plant’s adaptive responses.

While the importance of small heat shock proteins (sHSPs) in plant stress adaptation is well-documented, a significant gap remains in understanding the precise mechanisms underlying their action. To bridge this knowledge gap, we have embarked on a systematic exploration of the SmsHSP24.1 promoter region. In our previous paper, we have demonstrated the heat inducible chaperon activity of mitochondria localized SmsHSP24.1 protein and impact of transgenic overexpression on plant growth and development (Khatun et al., 2021). To further understand the molecular regulatory mechanism, we have subsequently generated a series of deletion constructs of SmsHSP24.1 promoter and fused with the uidA reporter gene (encoding beta-glucuronidase, GUS) to elucidate the putative cis-regulatory elements involved in abiotic stress signaling in the BARI begun-4 Eggplant variety. The SmsHSP24.1 promoter holds substantial promise not only for advancing our understanding of plant stress responses but also for its potential applications in crop genetic enhancement through transgenic techniques. This research underscores the significance of unraveling the intricacies of promoter-driven gene regulation and sets the stage for a deeper exploration of uncharted gene functions.

## 2. Materials and method

### 2.1. Plant materials and growth conditions

In this study, we used BARI Begun-4, an Eggplant (*Solanum melongena* L.) variety collected from the Bangladesh Agricultural Research Institute (BARI). The collected seeds were first surface sterilized in the laminar airflow with 1% Bavistin and 0.1% mercuric chloride (HgCl2) solution for a period of 5 minutes. Subsequently, the seeds underwent a series of four to five rinses using autoclaved distilled water. Afterward, to remove excess moisture, the seeds were carefully placed on blotting paper within the laminar airflow chamber. Once adequately dried, the surface-sterilized seeds were inoculated into jam jars containing solidified MS salt medium (Murashige & Skoog, 1962) supplemented with 3% sucrose to facilitate germination and seedling development. Following a two-week period under controlled conditions (16-hour photoperiod at 25.2°C), germinated seedlings were transferred to a hydroponic system utilizing Yoshida Solution (Yoshida et al., 2014). At three weeks of age, these seedlings were subjected to various stressors, including heat (45°C), salt (200 mM NaCl), and drought (150 mM mannitol) to better understand the SmsHSP24.1 expression in various tissues. Plants that were only treated with water were used as treatment controls (CT). Leaf samples were collected both before and after the stress treatment at 0, 1, 2, 4, 6, and 12 hours for the extraction of RNA and the subsequent investigation of gene expression.

### 2.2. Cloning and sequence analysis of SmsHSP24.1 promoter

The genomic DNA was extracted from 3-week-old Eggplant (*Solanum melongena* L.) young leaves using the CTAB method. The full-length upstream sequence, approximately 2 kilobase (kb) preceding the translation initiation codon (ATG) of small heat shock protein SmsHSP24.1 promoter (2000 bp to 1), was amplified by using a pair of PCR primers (HSP_Pro_1F/ HSP_Pro_1R; Supplementary Table 1). Subsequently, the amplified promoter sequence was cloned into the pCR-4-TOPO vector. To validate the sequence integrity, confirmation was carried out employing universal M13 reverse and forward primers-based Sanger sequencing. Putative conserved plant cis-acting regulatory elements and motifs within the SmsHSP24.1 promoter were analysed using different bioinformatic software i.e., PLACE (Higo et al., 1999) and plantCARE (Lescot et al., 2002).

### 2.3. Construction of SmsHSP24.1 Promoter: GUS expression vectors

The full-length (⁓2kb) SmsHSP24.1 promoter sequence was initially amplified and cloned into pCR-4-TOPO vector. Subsequently, three additional 5’ deleted fragments of varying lengths (1.5 kb, 1.0 kb, 0.5 kb) were also amplified using specific PCR primer pairs (HSP_Pro_2F, 3F, 4F/ HSP_Pro_2R, 3R, 4R; Supplementary Table 1).

After successful amplification, the four different fragments of SmsHSP24.1 promoter, like 2.0kb (−2000 bp to −1 bp from ATG of SmsHSP24.1 gene), 1.5kb (−1500 bp to −1 bp from ATG of SmsHSP24.1 gene), 1.0kb (−1000bp to −1 bp from ATG of SmsHSP24.1 gene) and 0.5kb (−500 bp to −1 bp from ATG of SmsHSP24.1 gene), were sub-cloned into entry vector (pL12R34-Ap) using *Sac*I and *Nco*I restriction sites. All these four 5′ deleted fragments were then fused with the uidA (beta-glucuronidase, GUS) reporter gene for the construction of SmsHSP24.1 promoter and uidA coding region along with Nopaline synthase (Nos) terminator expression vector (SmsHSP24.1+GUS+NosT). The GUS and Nos terminator regions were amplified from pCAMBIA1301 plasmid using the primer pair, GUS_F/NosT_R (Supplementary Table 1) and a high-fidelity DNA polymerase (KOD plus, Toyobo, Japan). The amplified GUS+Nos terminator portion was then cloned into the entry vector (pL12R34-Ap) containing the previously cloned four 5′ deleted SmsHSP24.1 promoter regions. Once the full expression cassette was confirmed with colony PCR, these complete expression cassette with four different deletions fragments of SmsHSP24.1 promoter were then transferred into plant expression vector pMDC100 having the NptII as selection marker gene in its backbone using LR recombinase mediated GatewayTM (Invitrogen, USA) cloning method (Figure: 1C).

**Figure 1:**
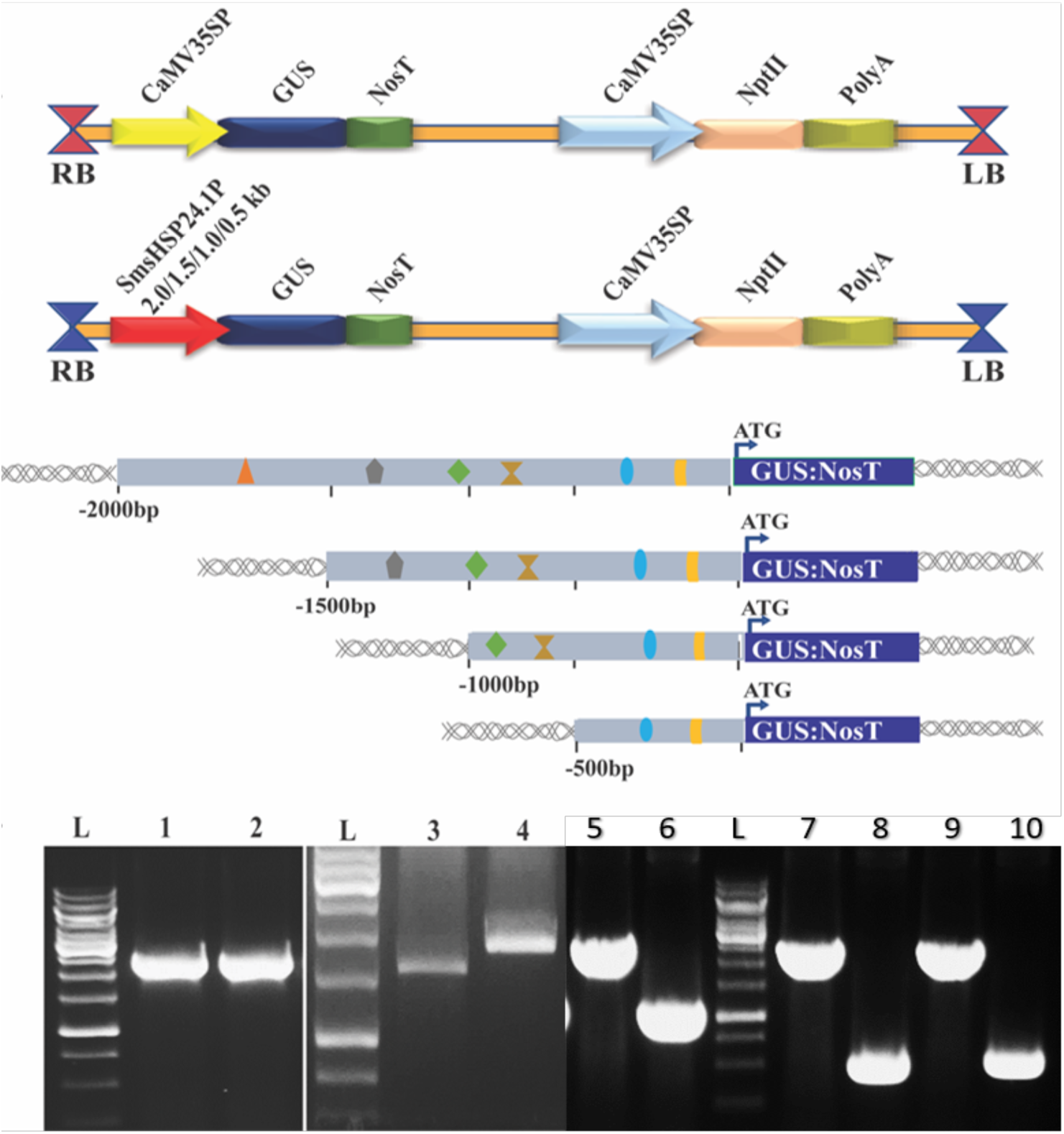
Construction of *SmsHSP24.1* and CaMV35S promoter driven *GUS* and *Nos* terminator (GUS:NosT) plant expression vectors used in this study. (A) A schematic representation of the CaMV35S promoter:GUS:NosT expression cassette. (B) A schematic representation of the *SmsHSP24.1* truncated promoter:GUS:NosT expression cassette and diagrammatic representation of four 5′ deletion variants of the *SmsHSP24.1* promoter:GUS:NosT expression cassette (C) PCR validation of the *SmsHSP24.1* promoter:GUS:NosT expression construct in *Agrobacterium* strain EHA105. Lane L shows a 1kb ladder (Thermo Scientific O’GeneRuler 1 kb DNA Ladder, ready-to-use #SM1163). Lanes 1, 3, 6, and 8 display amplified regions of the *SmsHSP24.1* promoter at sizes of 2.0kb, 1.5kb, 1.0kb, and 0.5kb, respectively. Lanes 2, 4, 5, 7, and 9 show the amplified 2.0kb GUS fragment. Lane 10 demonstrates the amplified CaMV 35S promoter region.

Additionally, for the construction of the CaMV35S promoter:GUS expression vector, the PCR-amplified GUS and Nos terminator from pCAMBIA1301 were directly cloned into the modified Entry clone 1 (pL12R34-Ap) under the CaMV35S promoter and Nos terminator between *Nco*I and *Xho*I restriction sites. And upon confirmation through PCR (Figure: 1C) transferred into plant expression vector pMDC100 following the same GatewayTM (Invitrogen, USA) cloning method. Finally, all these plant expression vectors were introduced indecently into the *Agrobacterium tumefaciens* EHA105 strain for transformation to the eggplant genome.

### 2.4. *Agrobacterium*-mediated genetic transformation of Eggplant

For *Agrobacterium*-mediated genetic transformation, we followed our previously published protocol (Khatun et al., 2022). Cotyledonary leaves of 21 days old *in vitro*-grown Eggplant seedlings were used. Each cotyledonary leaf was transversely cut into 2-3 pieces and subjected to infection with the EHA105 *Agrobacterium* strain harbouring the plant transformation constructs. We utilized our previously optimized protocol for BARI Begun-4 for stable transformation and regeneration. The transformed cells were selected on MS medium supplemented with 2.5 mg/l BAP (6-benzylaminopurine) and 100 mg/l kanamycin. Untransformed calli turned black and died at 100 mg/l kanamycin concentration. The micro-calli were then transferred to regeneration media for initiation of shoots. Following adequate elongation, the regenerated shootlets were transferred to rooting media. Elongated shoots-initiated root induction in full or half-strength MS medium supplemented with 1.0 mg/L IBA (Indole-3-butyric acid). After adequate root development, they were transferred into plastic pots. Finally, the acclimatized putative transgenic plants were transferred to greenhouse, where their growth and development were further monitored and studied (Figure 2A-F).

**Figure 2:**
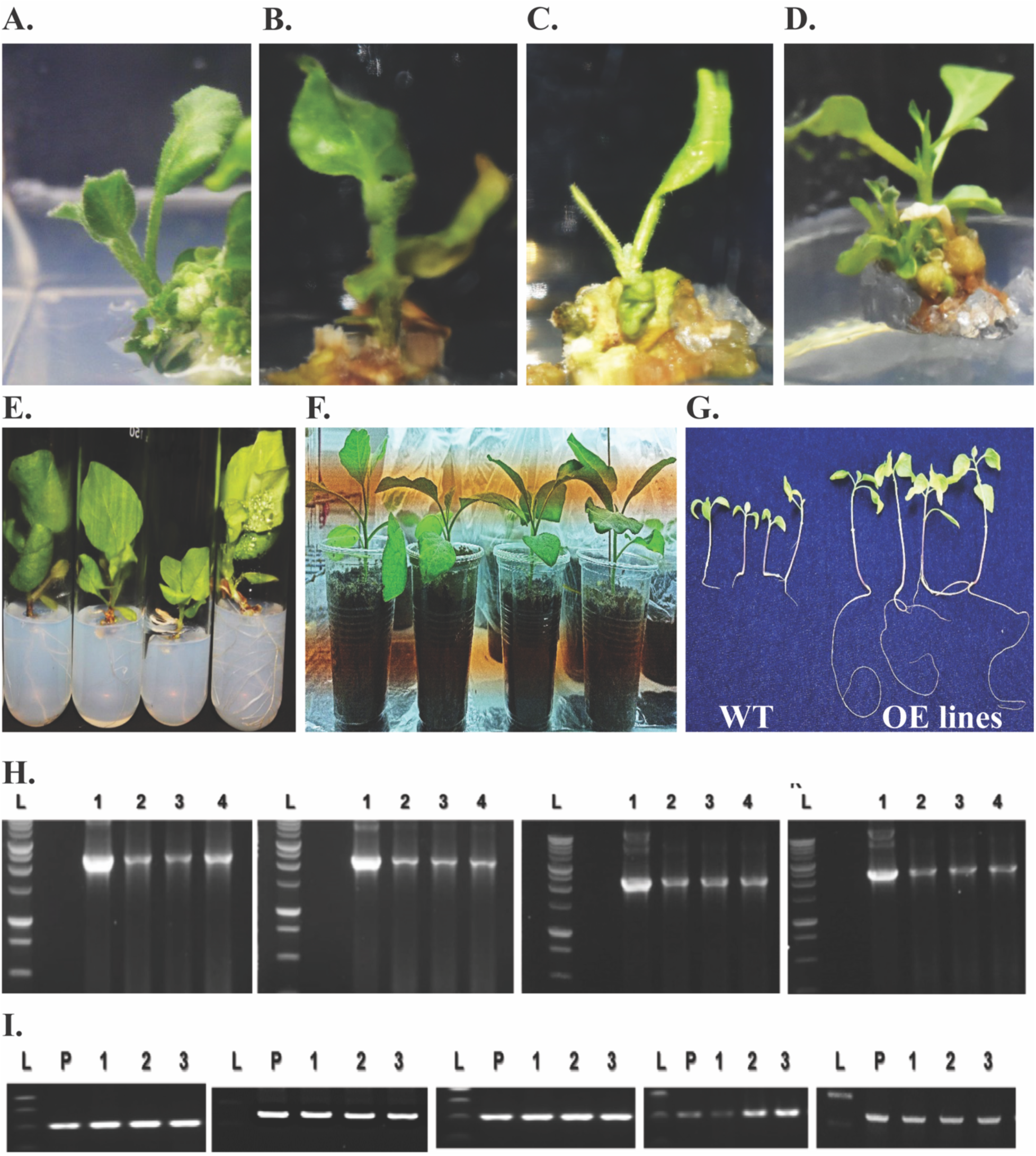
*Agrobacterium* mediated transformation of the *SmsHSP24.1* Promoter:GUS:NosT Expression Vector into the BARI Begun-4 Eggplant Variety. (A-D) Show the shoot regeneration of four *SmsHSP24.1* promoter:GUS:NosT transformed explants with deletions of 2.0kb, 1.5kb, 1.0kb, and 0.5kb. (E) Illustrates the *in vitro* rooting of four lines with 5′ deletions in the *SmsHSP24.1* promoter:GUS:NosT. (F) Depicts the acclimatization process of four distinct putative transgenic lines. (G) Highlights the selection of transgenic lines under a kanamycin concentration of 150 mg/l. (H) Display PCR amplification of the Gus+NosT terminator from transgenic eggplant lines with 5′ deletions in the *SmsHSP24.1* promoter:GUS:NosT, where Lane L is a 1kb ladder, Lane 1 is the Gus+NosT amplified region as a positive control, and Lanes 2-5 show the Gus+NosT amplified regions from the four transgenic lines (2.0kb, 1.5kb, 1.0kb, 0.5kb, respectively). (I) Shows PCR amplification of kanamycin resistance from lines transformed with either the *SmsHSP24.1* promoter:GUS:NosT or the CaMV35S promoter:GUS expression vectors.

### 2.5. Molecular screening and identification of transgenic Eggplant

Genomic DNA was extracted from all the putative T0 transgenic lines developed with five independent expression vectors, i.e., pMDC100:2.0 kbSmsHSP24.1 promoter+Gus+NosT, pMDC100:1.5 kbSmsHSP24.1 promoter+Gus+NosT, pMDC100:1.0 kbSmsHSP24.1 promoter+Gus+NosT, pMDC100:0.5 kbSmsHSP24.1 promoter+Gus+NosT and CaMV35S promoter+Gus+NosT along with control eggplant leaves using PureLinkgenomic plant DNA purification kit (Invitrogen, USA) according to the manufacturer’s instructions. To confirm transgene insertion, PCR screening was carried out to trace the presence of nptII gene sequence in the respective T-DNAs with the primer pair (NptII_F/NptII_R) (Supplementary Table 1) which produced an amplicon of ∼0.45 kb amplicon. Further confirmation was carried out using Gus+NosT primer pair produced ∼2.0 kb band length for the respective constructs (Figure: 2 H-I).

### 2.6. Acclimatization and generation of T3 transgenic Eggplants

The transgenic T1 seeds were collected from distinct PCR-positive T0 Eggplant lines. The collected seeds T0 along with wild type (WT) were first surface sterilized in the sample process describe above section (Plant materials and growth) These surface-sterilized seeds were later inoculated into jam jars containing solidified MS salt medium supplemented with 150 mg/L of kanamycin for selection of T1 transgenic lines for respective constructs. Non-transformed T1 seedlings stop root formation at 150 mg/l kanamycin concentration like WT seedlings (Figure: 2G). After 20 days of culture the healthy and well rooted seedlings were selected and transferred into the soil pot. For the generation of T2 and T3 transgenic seeds, the same procedure was followed as describe above. Finally, the T3 seedlings were used for subsequent SmsHSP24.1 promoter+GUS expression analysis.

### 2.7. Histochemical GUS staining

GUS (β-glucuronidase) assay was conducted using WT and T3 transformed eggplant lines according to the description of (Jefferson et al., 1987). Samples were selected from 5′ deleted four SmsHSP24.1 promoter and CaMV 35S promoter transformed transgenic as well as control eggplant lines with and without abiotic stress. The staining procedure involved the application of GUS assay buffer solution (composition:1 mg/ml X-Gluc, 100 µM Na2-EDTA, 100 mM sodium phosphate buffer pH 6.8, 0.5 mM potassium ferricyanide, 0.5 mM potassium ferrocyanide) followed by vacuum treatment with approximately 10 min with subsequent overnight incubation at 37 °C. After staining, samples were repeatedly washed with 70% (v/v) ethanol for removing the chlorophyll and then viewed under a microscope (Zeiss SteREO Discovery.V12, Germany). For each construct, samples were collected from at least three different transgenic lines.

### 2.8. qPCR expression analysis of GUS

Homozygous T3 generation of four different 5′ deletion of SmsHSP24.1 promoter transgenic lines along with CaMV 35S driven transgenic lines as well as WT seedlings were selected for exposure to various abiotic stress conditions. 3-week-old transgenic and WT seedlings were treated with salt stress (200 mM NaCl), heat stress (45°C, for 2 hr), and drought stress (150 mM mannitol) to investigate the transcript abundance of GUS and SmsHSP24.1 using quantitative RT-PCR analysis. The expression levels of GUS and SmsHSP24.1 were normalized to the housekeeping gene 18s rRNA, and the results were presented as expression fold changes relative to untreated control plants. Plants grown only with water served as untreated controls.

### 2.9. Total RNA isolation for RT-qPCR analysis

Total RNA was extracted from eggplant tissues using the PureLink Plant RNA Reagent kit (Invitrogen; https://www.thermofisher.com). RNA was treated with RNase-free DNase (NEB, http://www.neb.com) and used for first-strand cDNAs synthesis according to Verso cDNA Synthesis Kit (ThermoScientific; https://www.thermofisher.com). Real-time qRT-PCR was performed with SYBR™ Green PCR Master Mix and Applied Biosystems™ 7500 Real-Time PCR Systems (ThermoScientific; https://www.thermofisher.com). Eggplant 18S rRNA specific primer pair was used as an internal reference gene. All primer pairs used for Real-time qRT-PCR are mentioned in Supplementary Table 1.

### 2.10. Statical analysis

Statistical analyses were performed by one-way ANOVA and the differences between means were compared with Tukey HSD 0.5 values obtained for the particular dataset.

## 3. Results

### 3.1. Cloning of novel *SmsHSP24.1* promoter sequence from Eggplant (*Solanum melongena* L.)

In our previous study, we demonstrated a novel mitochondrion localized SmsHSP24.1 protein from Eggplant and characterized this protein under both normal and stressful circumstances (Khatun et al., 2021). In the transgenic overexpression lines, we observed rapid germination and seedling vigour compared to wild types along with improved abiotic stress resistance, particularly in temperatures up to 45 ^°^C in the field conditions. We therefore attempted to functionally characterize the *SmsHSP24.1* promoter to fully comprehend the signalling and expression regulation of the SmsHSP24.1 protein. In this study, first, we amplified the ⁓2kb upstream sequence (predicted as a small heat shock protein promoter) of the translation initiation codon (ATG) of small heat shock protein promoter (−2000 bp to −1 bp from SmsHSP24.1 ORF) by using a primer sets (HSP_Pro_1F/ HSP_Pro_1R, Supplementary Table 1) from genomic DNA of Eggplant (*Solanum melongena* L.) variety BARI begun-4. After the successful amplification of the full-length SmsHSP24.1 promoter sequence, we cloned the amplified product into the pCR-4-TOPO vector. Sequence confirmation was subsequently done by universal M13 reverse and forward primers. We further deposited the confirm full length promoter sequence in NCBI Gene bank (GenBank accession number: BankIt2754018/OR703026). The function of the cis-acting elements and the stress-specific core functional regions associated with this promoter was identified by deletion experiments.

### 3.2. Construction of *SmsHSP24.1* promoter:GUS+NosT deletion expression vector

After successfully cloning and confirming the pCR-4-TOPO vector having 2kb upstream sequence previously characterised as a small heat shock protein promoter, then three others 5′ deleted fragments of different lengths (1.5kb, 1.0kb, and 0.5kb respectively) were also amplified by using a pair of PCR primers (HSP_Pro_2F, 3F, 4F/ HSP_Pro_2R,3R,4R) (Supplementary Table 1). All four 5′ deleted fragments of *SmsHSP24.1* promoter (2.0kb, 1.5kb 1.0kb and 0.5kb) were then sub-cloned into entry vector (pL12R34-Ap) using *Sac*I and *Nco*I restriction sites. Followed by fused with the GUS reporter gene (GUS F/NosT R) amplified from pCAMBIA1301 plasmid and then cloned under *Nco*I and *Xho*I restriction sites for the construction of *SmsHSP24.1* promoter+GUS+NosT expression vectors. Furthermore, For the construction of CaMV35S promoter:GUS expression vector, PCR amplified GUS and Nos terminator from pCAMBIA1301 was directly cloned into the entry vector 1 (pL12R34-Ap). under CaMV35S promoter using the restriction enzymes *Nco*I and *Xho*I. All entry vectors then subsequently transferred into destination vector pMDC100. The backbone of pMDC100 vector already has plant selection marker gene kanamycin thus making it suitable for Eggplant transformation. LR recombinase mediated Gateway^TM^ (Invitrogen, USA) cloning strategy was used to make all expression vectors. Finally, *Agrobacterium tumefaciens* EHA105 strain was used as a host cell for the transfer of this GUS expression vector into eggplant (Figure. 2A-F).

### 3.3. Generation of *SmsHSP24.1* promoter+GusNosT stable transgenic Eggplant lines

We used BARI Begun-4 Eggplant variety for *Agrobacterium* mediated stable transformation. The four 5’ deletion *SmsHSP24.1* promoter:GUSNosT and CaMV35S promoter:GUS gene expression cassettes harbouring vectors were transformed into BARI Begun-4 Eggplant variety through our previously optimized *Agrobacterium*-mediated transformation and regeneration protocol (Khatun et al., 2021). After successful transformation, we generated four different 5′ deleted *SmsHSP24.1* promoter::GUSNosT fused transgenic eggplants lines along with CaMV35S promoter:GUSNosT fused lines. PCR confirmation with two pairs of primers (kanamycin primer sets produced ∼0.45 kb band length and Gus+NosT primer pair produced ∼2.0 kb band length) confirmed the T0 positive plants from all transgenic Eggplant lines (Figure: 2H-I). Homozygous T2 generation transgenic lines were selected under 150 mg/l kanamycin selection pressure and reconfirmed by PCR screening was selected for giving all abiotic stresses.

### 3.4. *SmsHSP24.1* promoter sequence analysis reveals various stress-responsive *cis*-regulatory elements indicating SmsHSP24.1 protein regulation under abiotic stresses

Sequence homology search of our newly cloned full-length promoter sequence (1957 kb upstream sequence of SmsHSP24.1 protein) from genomic DNA of BARI begun-4 eggplant variety against NCBI Eggplant Genome Database (Wei et al., 2020) (mitochondrial small heat shock protein, accession: E08). Further, we used Clustal Omega software (Madeira et al., 2022) for the sequence alignment of *SmsHSP24.1* promoter with NCBI reference accession E08.

We observed a total of 24 mismatches and 3 gaps in *SmsHSP24.1* promoter sequences. These mismatches could be due to different inbred lines (BARI begun-4 Bangladeshi Eggplant variety) used in this study. A notable change was spotted in the *SmsHSP24.1* promoter’s W-Box motif, where a cytosine takes the place of a thymine at the 817(+) position from the 5’ end. The CCAAT box motif in the *SmsHSP24.1* promoter has a cytosine instead of the thymine found in the E08 HSP promoter sequence at the 429(+) position from the 5’ end (Supplementary Figure 1).

To find cis-regulatory elements in the full-length *SmsHSP24.1* promoter sequence, we used three online tools: New PLACE (Higo et al., 1999), Plantpan 2.0 (Chow et al., 2016), and PlantCARE (Lescot et al., 2002), following their default settings. The results are shown in Figure 3. We identified multiple TATA Box and CAAT box motifs in the *SmsHSP24.1* promoter, detailed in Supplementary Table 2. Using TSSPlant software (www.softberry.com/berry.phtml?topic=tssplant&group=programs&subgroup=promoter), we predicted five potential transcription start sites (TSS). The TSS closest to the start codon was selected and is highlighted in Figure 3.

**Figure 3:**
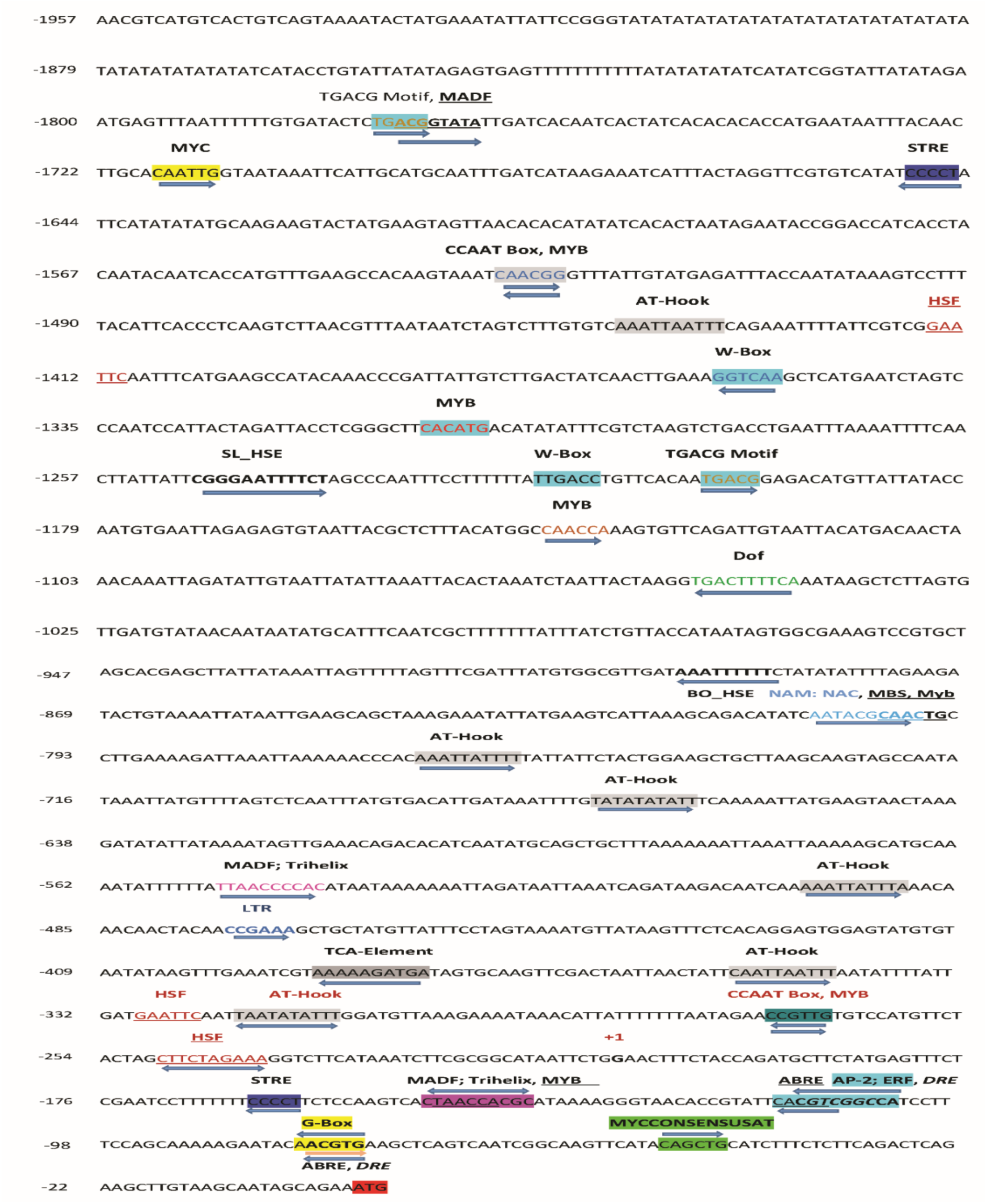
Visualization of *cis*-regulatory motifs in the *SmsHSP24.1* promoter. The image depicts various *cis*-regulatory motifs highlighted in different colors, located upstream of the translation start site on the *SmsHSP24.1* promoter sequence. Each color represents a distinct type of cis-regulatory element, illustrating their spatial arrangement.

Our analysis predicts that the full-length promoter contains various stress-responsive cis-regulatory elements. These include the LTR (low-temperature response), TCA (salicylic acid response), ABRE (ABA response), G-BOX (light response), DRE core (dehydration response), TGACG motif (MeJA response), CCAAT Box (drought response), STRE (stress regulator), MYCCONSENSUSAT, and MADF: trihelix binding motif (stress and pathogen response). In addition to these motifs, the *SmsHSP24.1* promoter harbors several transcription factor core sites known to be involved in stress-responsive plant growth. For instance, the AT-Hook motif, implicated in flowering initiation, developmental processes, and stress response, appears 154 times. Other transcription factors, including AP2; ERF (hormone and abiotic stress regulation), NAM, NAC (salt stress tolerance regulation), HSF (heat stress response), MYB transcription factor recognition site (transcriptional regulation, drought response), and WRKY proteins (which bind in W-Box motifs to regulate plant development and stress responses), are also present in the *SmsHSP24.1* promoter.

In the analysis of the *SmsHSP24.1* promoter, the shortest deletion fragment of 0.5 kb was found to contain a diverse array of cis-regulatory motifs indicative of its responsiveness to various stress conditions. This fragment encompasses three motifs associated with heat stress, two with cold stress, three with salt stress, one specific to drought stress, and four motifs related to a multifunctional stress response. Specifically, under heat stress conditions, this promoter fragment becomes active through one STRE and two HSF motifs located at positions +47 (−), −122(+), and −42(+) respectively, which are known to facilitate HSP protein expression. This fragment also contains two number of salt tress related motifs naming AP2; ERF with position +96(+), ABRE with position +94(−), +129(−).

In case of 1.0 kb deletion fragment, in addition to the motifs found in the 0.5 kb fragment, also consist of the BO_HSE motif for heat stress (−688bp), NAM and NAC motifs for salt stress tolerance (−599bp), MBS for drought inducibility (−593bp), and MADF (−344bp) a trihelix motif for salt and drought stress responsiveness. These additions enhance the fragment’s capability to initiate stress tolerance mechanisms.

The 1.5 kb fragment retains all motifs identified in the shorter fragments and introduces several additional HSE/HSF motifs (at positioned −1039, −1042 and −1209 respectively) indicating a strong inclination towards heat stress responsiveness. Interestingly, this fragment lacks motifs for cold and salt stress found in smaller fragments but includes MYB (−934), DOF (−844), TGACG Motif (−996), and W-box (−937 and −1151), suggesting a complex network of stress response beyond heat stress.

The largest fragment analysed, the 2.0 kb segment, presumed to be the full-length promoter, encompasses all previously identified motifs across the fragments and introduces additional elements like STRE, CCAAT box, MADF, and TGACG motif, further amplifying its response to heat and various abiotic stresses. This comprehensive in silico analysis demonstrates the *SmsHSP24.1* promoter’s intricate regulation of HSP protein expression under heat, cold, salt, and drought stresses, as well as its role in biotic stress and growth promotion (Table: 1).

**Table 1:**
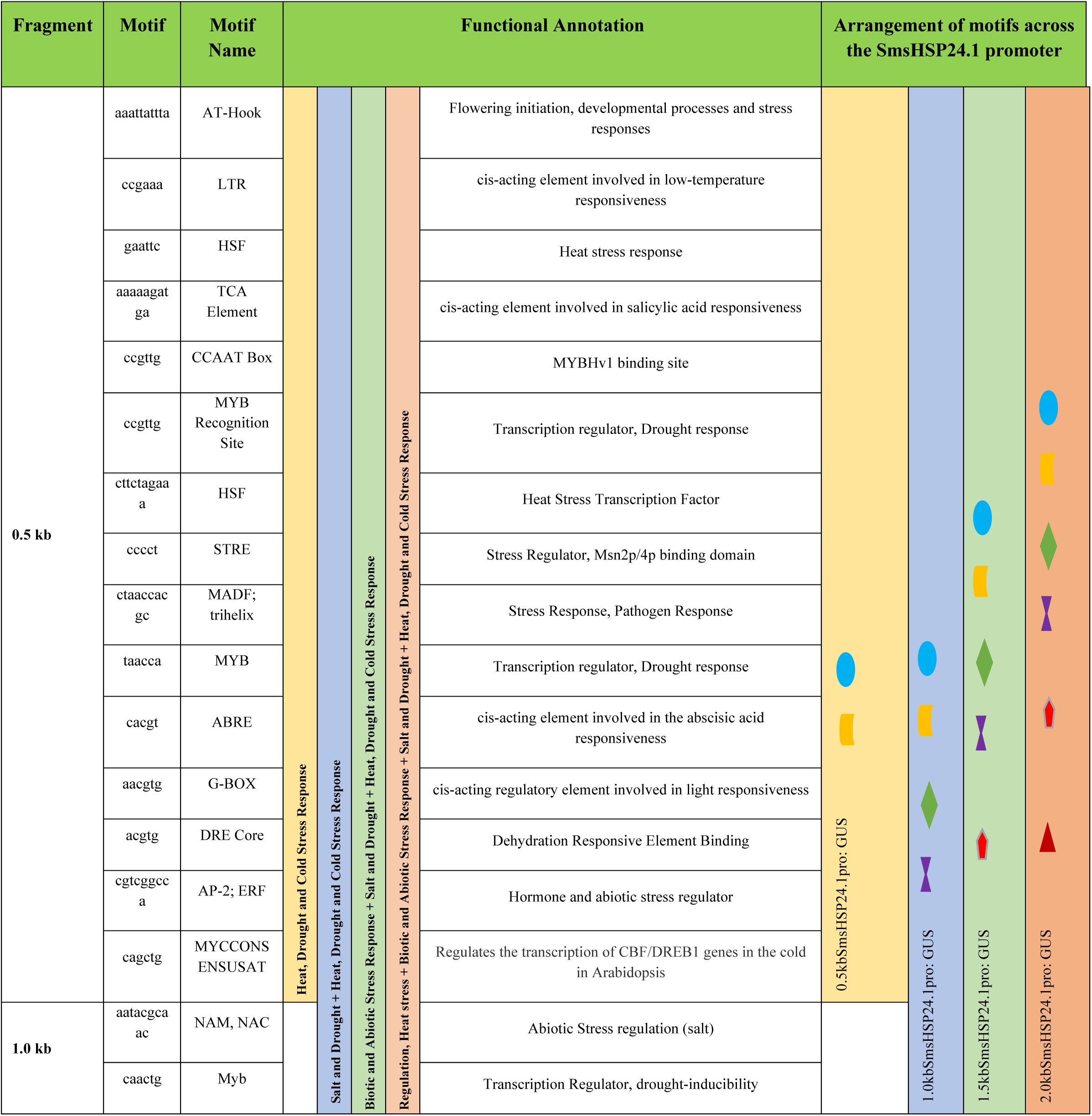
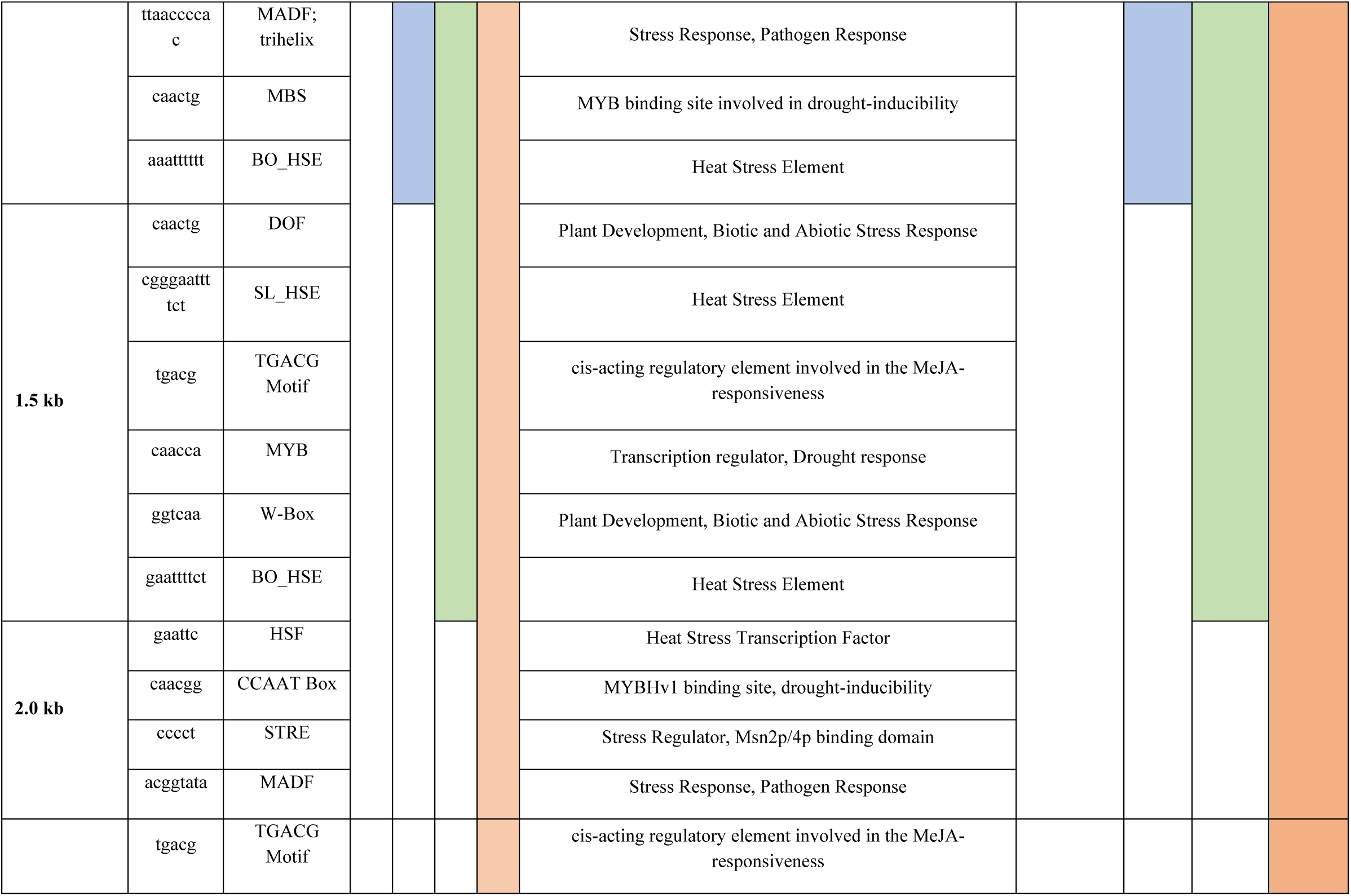
Predicted stress-responsive *cis*-regulatory elements in the ***SmsHSP24.1* Promoter**.

### 3.5. Histochemical analysis of 5′-truncated *SmsHSP24.1* promoter revealed the inducible function under abiotic stresses

We performed GUS histochemical staining assays on the transgenic eggplants lines overexpressing different 5′ deletion constructs of the *SmsHSP24.1* promoter:GUSNosT and CaMV35spromoter: GUSNosT expression cassettes. Homozygous T2 progeny, selected based on resistance to 150 mg/l kanamycin and subsequently verified through PCR, were subjected to various abiotic stresses to elucidate the functional cis-regulatory motifs within the *SmsHSP24.1* promoter. Three-week-old Eggplant (Solanum melongena L.) seedlings were exposed to a range of stress conditions, including 45°C to simulate heat stress, 150 mM mannitol for osmotic or drought stress, and 200 mM sodium chloride (NaCl) to induce salt stress, with results documented in Figures 4.

**Figure 4:**
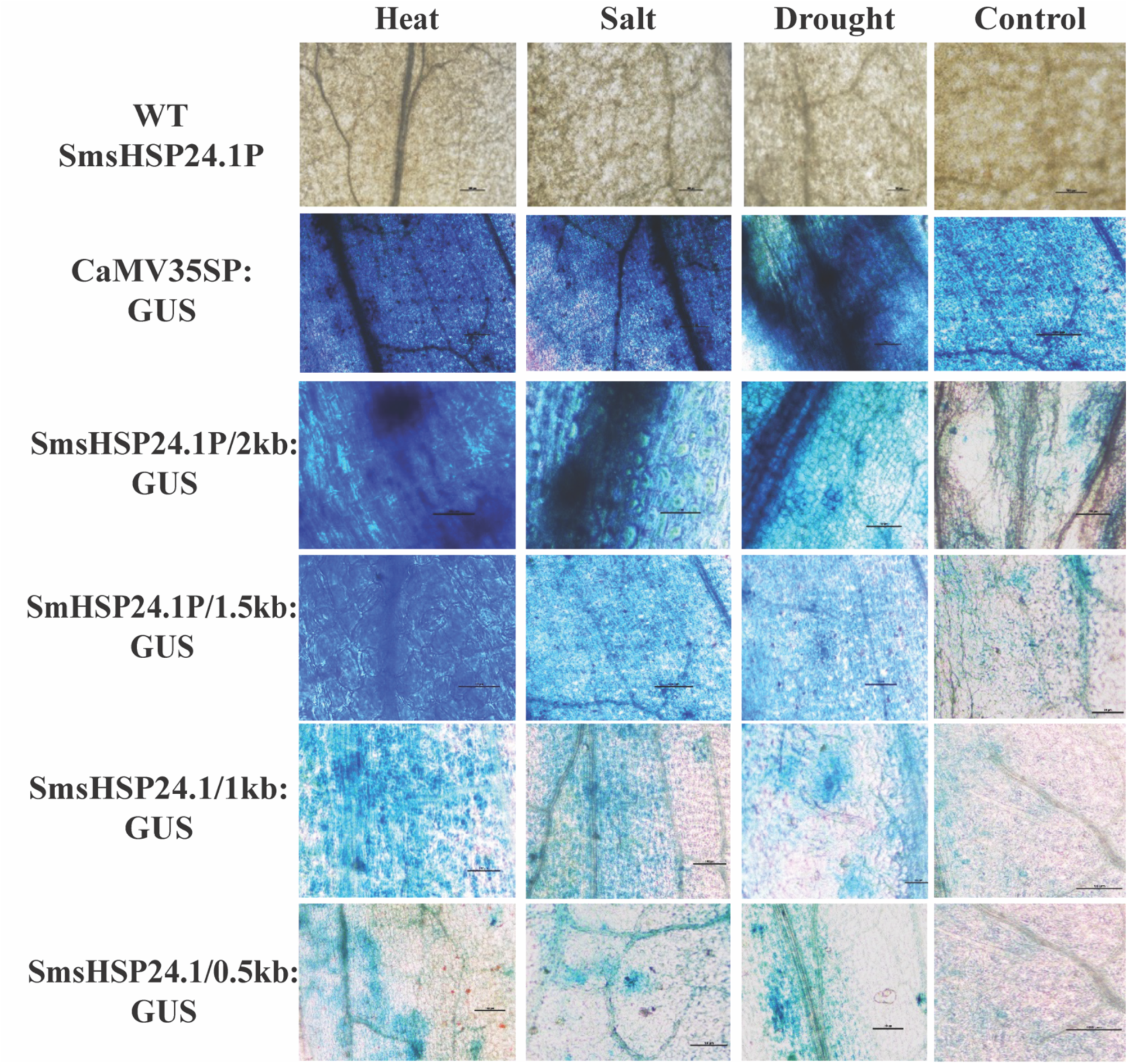
Histochemical staining of transgenic Eggplants exposed to different abiotic stresses such as heat (45 ^ο^C), salt (250 mM NaCl), and drought (150 mM Mannitol). This figure illustrates the *GUS* expression in leaf tissues of transgenic eggplants carrying the full-length 2.0kb *SmsHSP24.1* promoter:GUS:NosT and the CaMV35S promoter+GUS:NosT expression cassettes under various abiotic stress conditions, including heat, salt, and drought. Notably, strong GUS expression was observed under these stress conditions, with a modest presence in all transgenic lines in a controlled environment. Conversely, GUS activity was absent in the wild-type (WT) plants under both control and stress conditions. Remarkably, the most intense GUS expression was recorded in leaves with the 2.0kb *SmsHSP24.1* promoter subjected to heat stress.

The objective of histochemical staining was to quantitatively assess the expression of the GUS reporter gene across the different transgenic eggplant lines under these stress conditions. As expected, seedlings bearing the intact 2.0 kb *SmsHSP24.1* promoter-GUS construct displayed robust GUS expression under heat stress, thereby confirming the inducible capability of this promoter segment. Conversely, lines with 5′ deletions of the *SmsHSP24.1* promoter exhibited reduced GUS expression when challenged with heat, drought, and salt stresses, indicating a gradational decline in stress-responsive promoter activity. In comparison, transgenic lines harboring the CaMV35S promoter-GUS cassette showed consistent GUS expression independent of the stress condition applied, demonstrating the constitutive expression profile of the CaMV35S promoter. No GUS activity was detected in the leaf tissues of wild-type plants, as illustrated in WT SmsHSP24.1P in Figure 4. This differential expression pattern underlines the critical regulatory elements within the *SmsHSP24.1* promoter that mediate stress-induced gene expression, offering insights into the molecular mechanisms governing plant stress responses.

Heat stress imposed 21 days old transgenic Eggplant seedlings harbouring full-length 2.0kb *SmsHSP24.1* promoter:GUS:NosT expression cassette showed strong GUS expression and revealed the function of this promoter in an inducible way, whereas GUS expression was gradually declined in the 5′ deleted fragment of *SmsHSP24.1* promoter harbouring transgenic plants under heat, salt or even drought stress condition. The histochemical staining of transgenic leaf tissues harbouring CaMV35spromoter+GUSNosT expression cassette showed strong GUS expression irrespective of heat, salt, and drought stress conditions while GUS activity was not detected in leaf tissues of the WT plants (Figure 4).

### 3.6. qPCR analysis of 5′-truncated *SmsHSP24.1* promoter reveals significant gene expression in response to abiotic stresses

Further, expression levels of GUS transcripts fused with 5’ truncated *SmsHSP24.1* promoter was validated through time-dependent quantitative RT-PCR analysis, using RNA isolated from the leaves of 3-week-old transgenic and wild type Eggplant (*Solanum melongena* L.) seedlings. These seedlings were subjected to stress treatments of 200 mM NaCl for salt stress, 45°C for heat stress, and 150 mM mannitol for drought stress. The study encompassed the examination of expression patterns for both the native copy of *SmsHSP24.1* transcript and the GUS reporter gene driven by truncated *SmsHSP24.1* promoter. The housekeeping gene 18s rRNA was used to normalize the expression of the native *SmsHSP24.1* transcript and overexpressed 5’ deletion SmsHSP24.1: GUS constructs. The RT-PCR data were expressed as fold changes in expression relative to untreated seedlings, providing insights into the gene’s response under specific stress conditions, as depicted in Figure 5.

**Figure 5:**
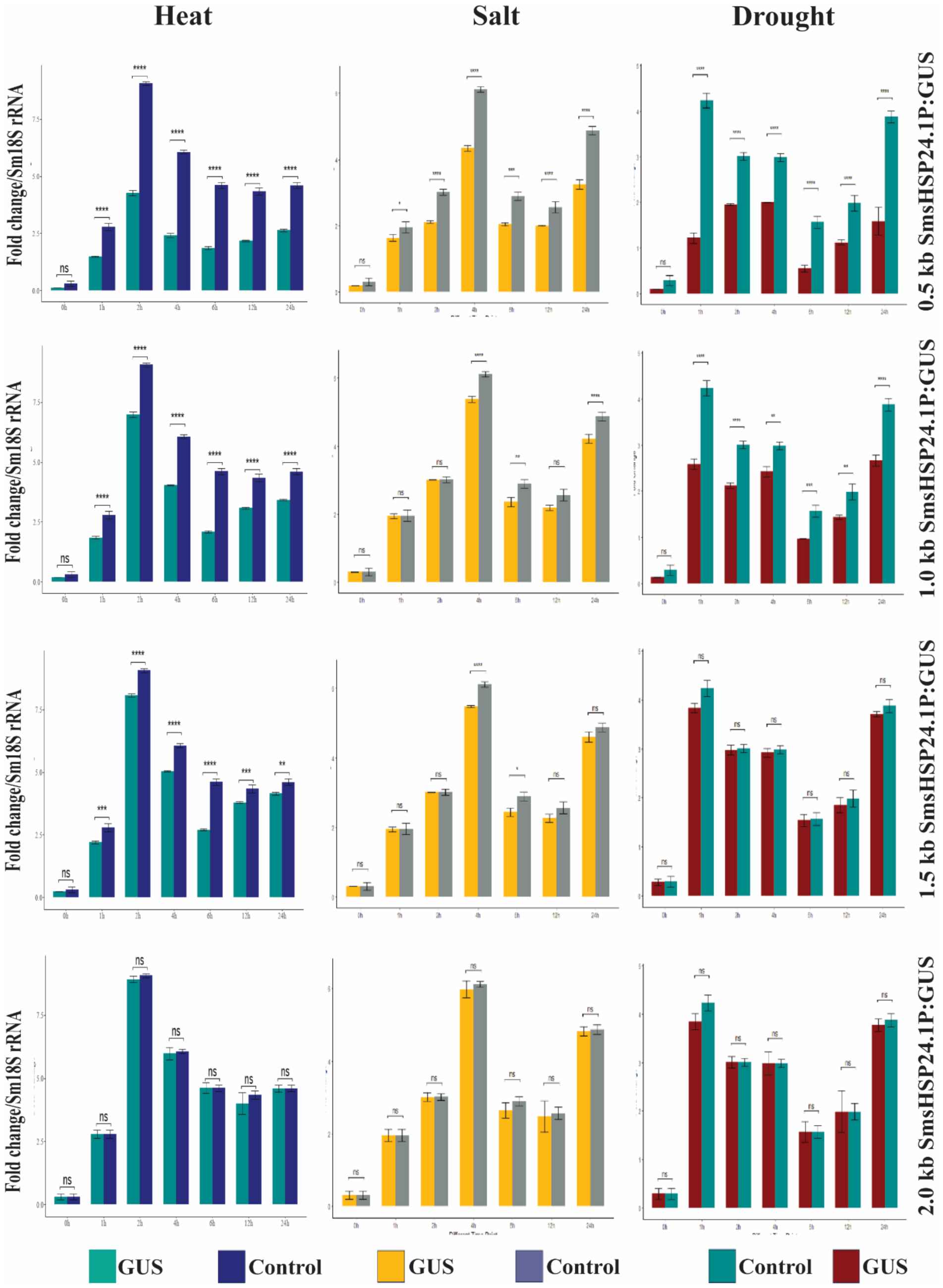
Time-dependent quantitative RT-PCR analysis under different abiotic stress conditions. This figure illustrates the responsiveness of the full-length *SmsHSP24.1* promoter to heat (45 ^ο^C), salt (250 mM NaCl), and drought (150 mM Mannitol) stresses over time periods of 0h, 1h, 2h, 4h, 6h, 12h and 24h respectively. The analysis reveals that the SmsHSP24.1 promoter exhibits a higher level of response to heat stress when compared to salt and drought conditions, indicating its significant role in the plant’s heat stress adaptation mechanism.

Our results revealed that heat stress elicited the most significant increase in relative fold change in gene expression, affecting both the native *SmsHSP24.1* gene and the GUS reporter gene under the control of the same *SmsHSP24.1* promoter. Following the initiation of heat stress, the native *SmsHSP24.1* transcripts surged, reaching an approximate 8-fold increase at the two-hour mark, before gradually diminishing. Similar to the native SmsHSP24.1 gene transcript, GUS reporter gene in plants equipped with the full-length 2.0kb *SmsHSP24.1* promoter-GUSNosT construct, exhibited a nearly 9-fold escalation in expression.

The expression dynamics of the native *SmsHSP24.1* transcript and overexpressed *GUS* transcript during salt stress were diverged notably from the heat stress. During salt stress, each transcript was highly upregulated at 4 hr of time point. At 4 hr of salt stress, an approximately 5-fold increase was observed compared to the control. The expression of the native *SmsHSP24.1* and *GUS* transcript were gradually reduced up to the 12 hr of salt stress but surged again at the 24 hr of salt stress (Figure. 5). In this case, the expression pattern was very much similar to 4 hr of salt stress.

When Eggplant seedlings were treated with mannitol as drought stress, both *GUS* and *SmsHSP24.1* transcripts exhibited rapid upregulation, achieving maximum expression at 2 hr post-treatment. This followed by a steady decrease across a 12-hr period. By the 24-hr exposure of drought stress, the expression had fallen to 1.5-fold lower than what was recorded 1 hr after the onset of stress (Figure 5).

## 4. Discussion

The regulation of plant growth, development, and adaptation through gene expression is intricate, heavily relying on the functionality of promoters and their cis-acting elements in fine-tuning transcription. With the advencement of transgenic and gene-editing technologies, the strategic selection and application of well-characterized promoters become fundamental. These promoters play critical role efficient genetic transformation vectors, elucidating gene functions, and enabling precise control over the expression of target genes in transgenic plant studies.

Experimental modulation of transgene expression typically involves linking the gene of interest to a promoter known for its specific expression patterns and intensities within the host organism. While universally strong promoters like those from the cauliflower mosaic virus (35S promoter) and the figwort mosaic virus (34S promoter), alongside plant constitutive gene promoters such as actin and ubiquitin, are favoured for their broad expression across various tissues (Potenza et al., 2004), their application is not without limitations. This underscores the utility of tissue-specific promoters which direct expression to specific cell types or tissues. Strong tissue-specific promoters that are expressed only in vascular tissues (Liu & Jia, 2003), anthers (Kato et al., 2010), or seed tissues are a few examples of those that have been utilized to drive plant transgenes (Furtado et al., 2008; Jones et al., 2005; Qu & Takaiwa, 2004).

A pivotal aspects of functional gene analysis in transgenic plants is the ability to precisely regulate transgene expression. This precision is crucial when constant up- or down-regulation of a target gene could negatively impact plant development, consume substantial resources, or when assessing the transgene’s effects across various conditions. Inducible promoters, which are activated by external stimuli such as heat, drought, or light, offer a refined control, enabling researchers to tailor gene expression to specific environmental cues or developmental stages (Argüello-Astorga & Herrera-Estrella, 1998; Corrado & Karali, 2009; Moore et al., 2006; Padidam et al., 2003). Although constitutive promoters are widely used, the specificity afforded by inducible and tissue-specific promoters significantly enhances the effectiveness of research by providing more controlled expression patterns and reducing unintended consequences of unregulated gene expression.

Our research delves into the stress-responsive behaviour of the SmsHSP24.1 protein, which has been previously shown to react sensitively under various environmental stresses such as heat, salt, and mannitol-induced drought conditions, highlighting a rapid increase in its transcript levels. This sensitivity underscores the protein’s role in stress response mechanisms, a notion further supported by the observed abiotic stress tolerance in BARI Begun-4 eggplants overexpressing *SmsHSP24.1* (Khatun et al., 2021).

To further understand the regulatory mechanisms governing *SmsHSP24.1* expression, we constructed and overexpressed four deletion constructs of its promoter fused with *GUS* in BARI Begun-4 eggplant variety. This approach illuminated the promoter’s role in stress and developmental regulation, as evidenced by the *GUS* gene expression pattern under different 5’ truncated *SmsHSP24.1* promoter. The inclusion of a CaMV35S promoter+GUSNosT expression cassette served as a benchmark for characterizing *GUS* expression levels in transgenic plants, revealing a gradual decrease in GUS activity from the 2.0kb *SmsHSP24.1* promoter:GUSNosT to the 0.5kb truncated construct. This pattern was particularly pronounced under heat stress conditions, where the full-length promoter demonstrated enhanced *GUS* expression, attributable to the presence of three HSF (Heat Stress Transcription Factor) binding motifs within the 2.0kb region (Kato et al., 2010; Liu & Jia, 2003).

The differential *GUS* expression patterns under salt, and drought stresses further elucidate the strategic placement of stress-responsive elements along the *SmsHSP24.1* promoter. The most comprehensive response was observed in the 2.0kb full length construct, most probably the full-length construct is equipped with major salt-responsive elements (NAM, NAC, Dof, W-box, and STRE) compared to notably reduce in *GUS* expression in the 0.5kb construct due to the absence of these key elements (Venter, 2007). The full length 2kb *SmsHSP24.1* promoter sequence has CCAAT box responsible for drought inducibility is absent in 1.5kb and 1.0 kb has Myb (responsible for drought inducibility) missing in 0.5kb. GUS expression is significantly decreased from 2.0kb to 0.5kb because of the loss of motifs that are required for promoter activation during stress.

Our findings suggest the full-length (∼2kb) *SmsHSP24.1* promoter’s potential as a versatile tool for crop improvement and gene function elucidation, given its array of biotic and abiotic stress-responsive elements as it is mentioned in the table 4.5. This study confirms the full-length promoter’s superior responsiveness to environmental stresses, supported by a robust set of regulatory motifs crucial for stress-induced gene expression (Argüello-Astorga & Herrera-Estrella, 1998; Corrado & Karali, 2009). All the motifs present in *SmsHSP24.1* promoter strongly justify its expression profile observed in GUS assay and signify its inducible expression in the transgenic line.

In conclusion, the *SmsHSP24.1* promoter exemplifies the critical balance between universal and specific regulatory elements, offering a promising avenue for enhancing plant resilience to environmental stresses through genetic engineering. Our work aligns with the growing body of literature advocating for the strategic use of well-characterized promoters in plant biotechnology research, paving the way for future advancements in agricultural genetic improvement.

---

**Supplementary Table 1:**
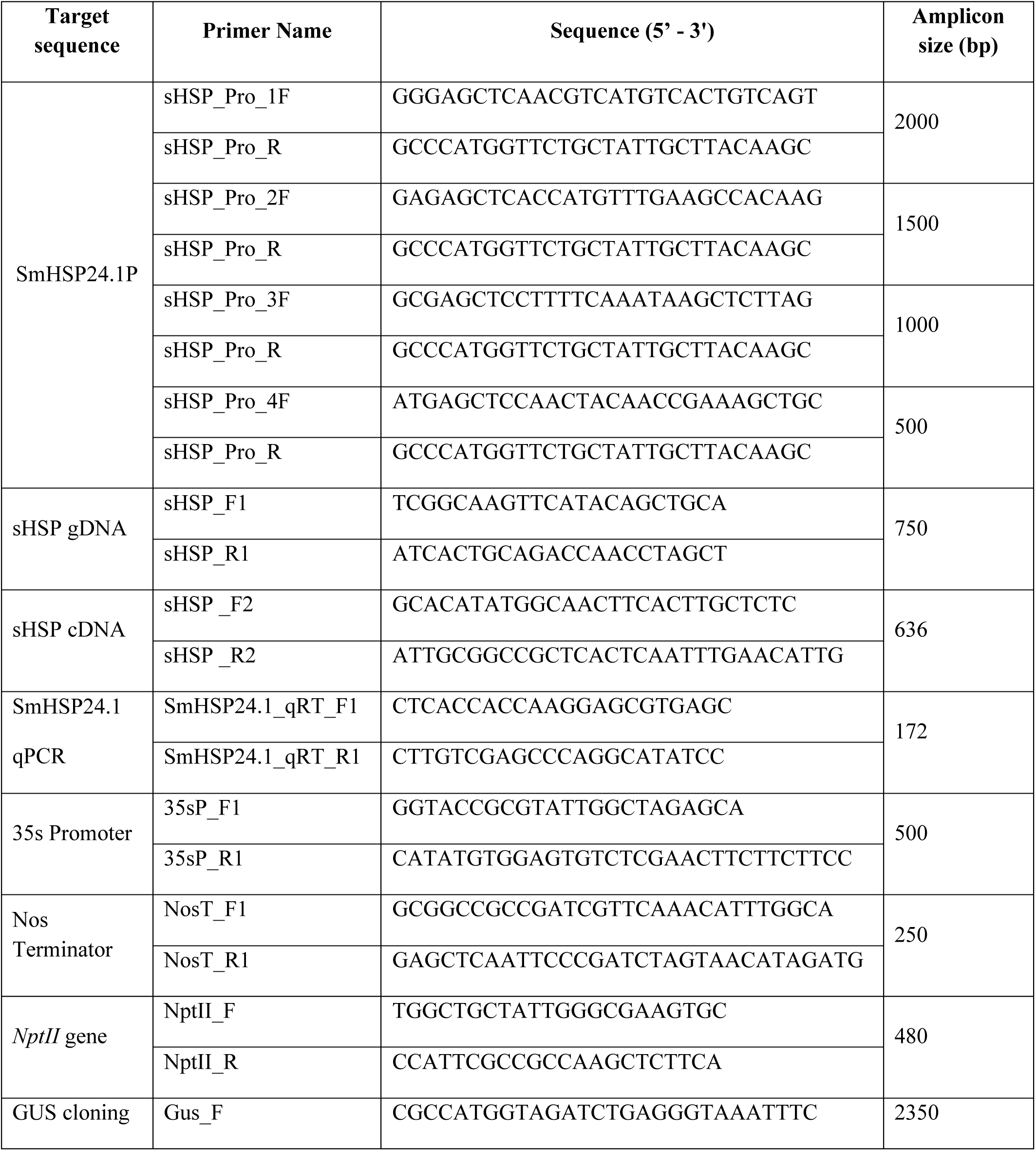
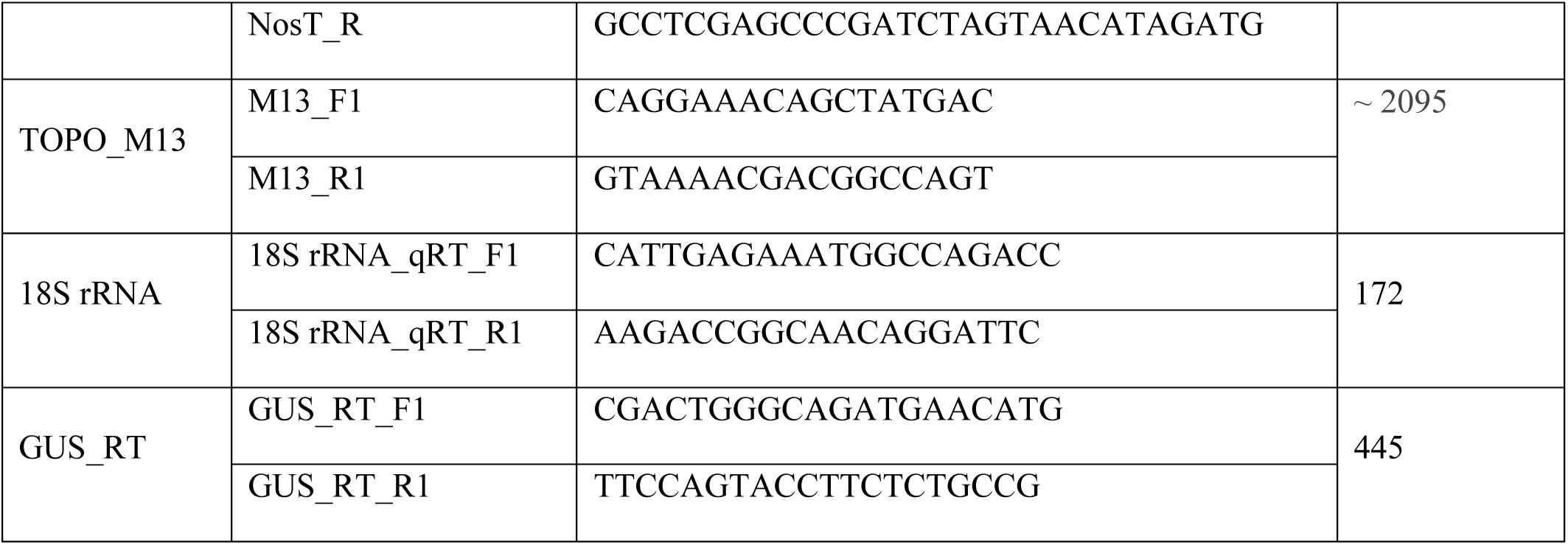
List of oligonucleotide primer sequences used in this study.

**Supplementary Table 2:** List of predicted TATA-box and CAAT-box in the *SmsHSP24.1* Promoter. The nearest TATA-box to the transcription start site (TSS), identified as TATABOX5 (sequence TTATTT), is located 75 bp upstream of the TSS. A CAAT-box (sequence CAAT) is found 31 bp upstream relative to TATABOX5, suggesting their roles in transcription initiation and regulation within the SmsHSP24.1 promoter.

| Motif Name | Motif Position and Direction (from 5' end of the Promoter) | Motif Sequence |
| --- | --- | --- |
| TATABOX2 | 1024 (+) | TATAAAT |
| TATABOX2 | 1240 (+) | TATAAAT |
| TATABOX3 | 1636 (-) | TATTAAT |
| TATABOX4 | 104 (-) | TATATAA |
| TATABOX4 | 127 (-) | TATATAA |
| TATABOX4 | 149 (-) | TATATAA |
| TATABOX5 | 971 (+) | TTATTT |
| TATABOX5 | 1196 (+) | TTATTT |
| TATABOX5 | 1660 (-) | TTATTT |
| TATABOX5 | 1669 (+) | TTATTT |
| CAATBOX1 | 192 (-) | CAAT |
| CAATBOX1 | 200 (+) | CAAT |
| CAATBOX1 | 241 (+) | CAAT |
| CAATBOX1 | 243 (-) | CAAT |
| CAATBOX1 | 258 (-) | CAAT |
| CAATBOX1 | 266 (+) | CAAT |
| CAATBOX1 | 452 (+) | CAAT |
| CAATBOX1 | 577 (-) | CAAT |
| CAATBOX1 | 624 (+) | CAAT |
| CAATBOX1 | 726 (+) | CAAT |
| CAATBOX1 | 753 (+) | CAAT |
| CAATBOX1 | 778 (+) | CAAT |
| CAATBOX1 | 834 (-) | CAAT |
| CAATBOX1 | 867 (-) | CAAT |
| CAATBOX1 | 944 (+) | CAAT |
| CAATBOX1 | 959 (+) | CAAT |
| CAATBOX1 | 1104 (-) | CAAT |
| CAATBOX1 | 1151 (+) | CAAT |
| CAATBOX1 | 1237 (+) | CAAT |
| CAATBOX1 | 1260 (+) | CAAT |
| CAATBOX1 | 1273 (-) | CAAT |
| CAATBOX1 | 1352 (+) | CAAT |
| CAATBOX1 | 1452 (+) | CAAT |
| CAATBOX1 | 1623 (-) | CAAT |
| CAATBOX1 | 1634 (+) | CAAT |
| CAATBOX1 | 1892 (+) | CAAT |
| CAATBOX1 | 1947 (+) | CAAT |

**Supplementary Figure 1:**
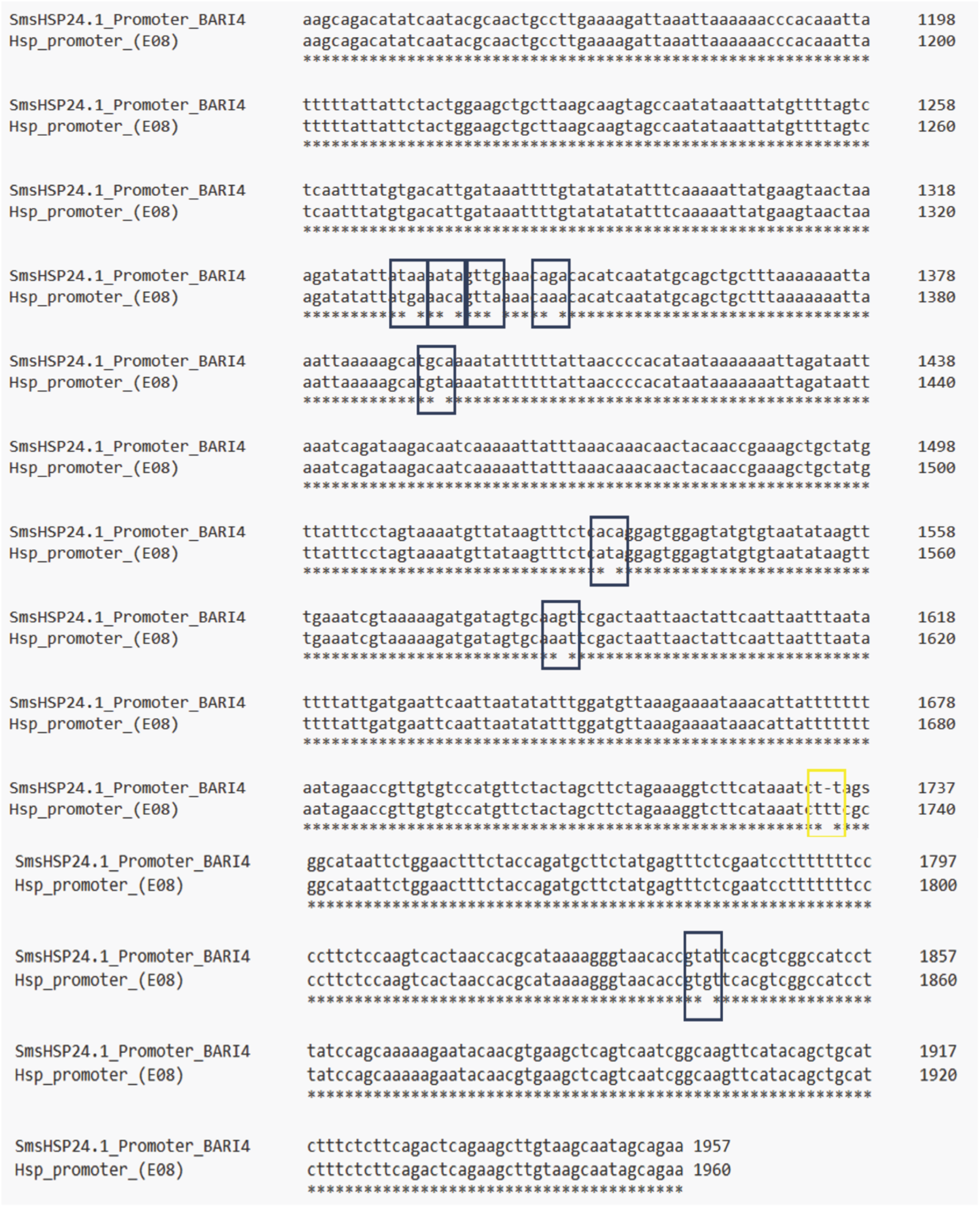
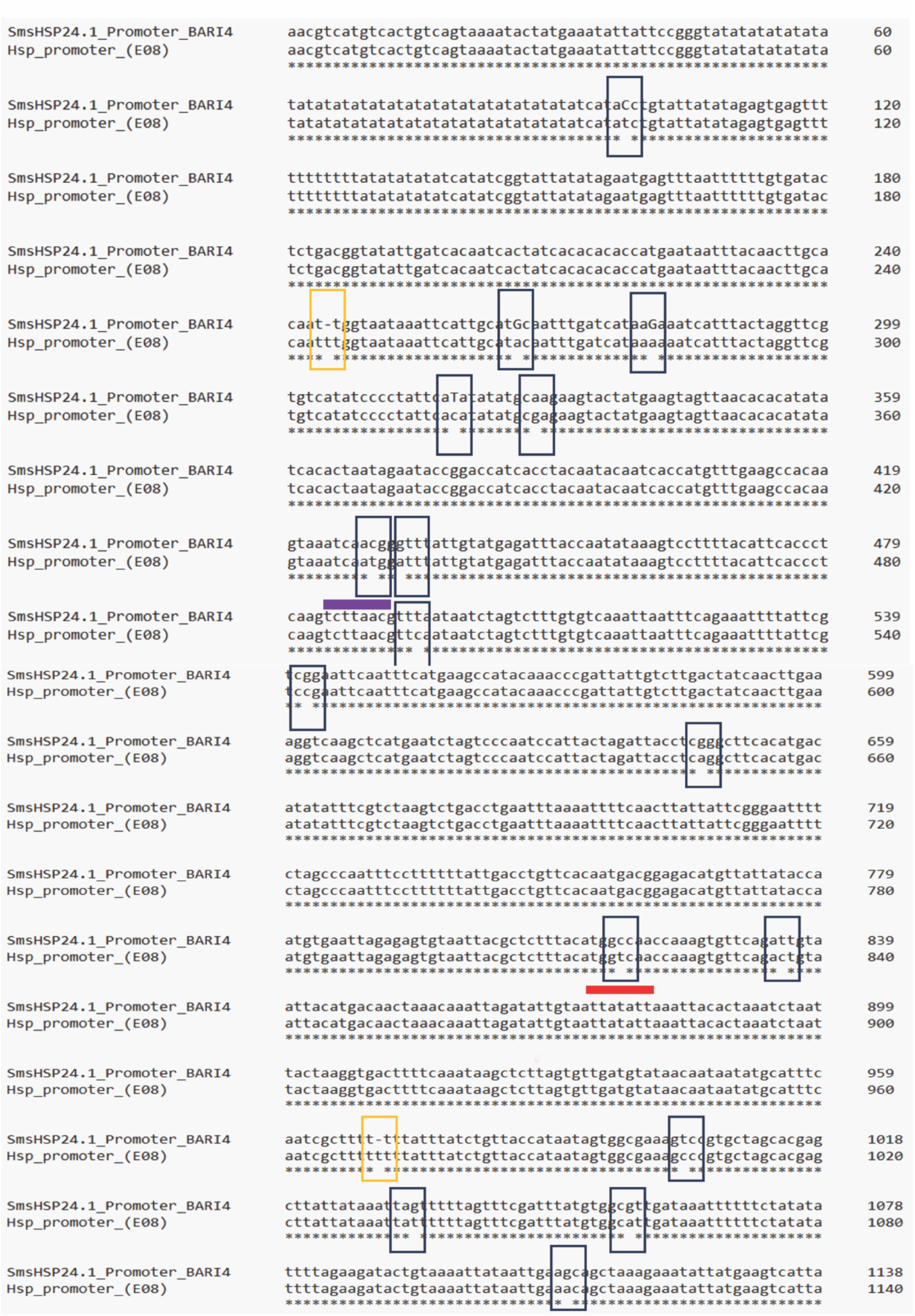
Sequence Alignment of *SmsHSP24.1* Promoter from BARI Begun 4 Eggplant. This figure presents a comparative sequence alignment between the full-length (2kb upstream of the gene) *SmsHSP24.1* promoter sequence obtained from the genomic DNA of the BARI Begun 4 eggplant variety and a draft mitochondrial small heat shock protein promoter sequence from a database. The alignment highlights similarities and differences, illustrating the degree of conservation and potential regulatory elements within the promoter region that may contribute to its function in heat shock response.

